# High fusion and cytopathy of SARS-CoV-2 variant B.1.640.1

**DOI:** 10.1101/2023.09.06.556548

**Authors:** William Bolland, Vincent Michel, Delphine Planas, Mathieu Hubert, Florence Guivel-Benhassine, Françoise Porrot, Isabelle Staropoli, Mélissa N’Debi, Christophe Rodriguez, Slim Fourati, Matthieu Prot, Cyril Planchais, Laurent Hocqueloux, Etienne Simon-Lorière, Hugo Mouquet, Thierry Prazuck, Jean-Michel Pawlotsky, Timothée Bruel, Olivier Schwartz, Julian Buchrieser

## Abstract

SARS-CoV-2 variants with undetermined properties have emerged intermittently throughout the COVID-19 pandemic. Some variants possess unique phenotypes and mutations which allow further characterization of viral evolution and spike functions. Around 1100 cases of the B.1.640.1 variant were reported in Africa and Europe between 2021 and 2022, before the expansion of Omicron. Here, we analyzed the biological properties of a B.1.640.1 isolate and its spike. Compared to the ancestral spike, B.1.640.1 carried 14 amino acid substitutions and deletions. B.1.640.1 escaped binding by some anti-NTD and -RBD monoclonal antibodies, and neutralization by sera from convalescent and vaccinated individuals. In cell lines, infection generated large syncytia and a high cytopathic effect. In primary airway cells, B.1.640.1 replicated less than Omicron BA.1 and triggered more syncytia and cell death than other variants. The B.1.640.1 spike was highly fusogenic when expressed alone. This was mediated by two poorly characterized and infrequent mutations located in the spike S2 domain, T859N and D936H. Altogether, our results highlight the cytopathy of a hyper-fusogenic SARS-CoV-2 variant, supplanted upon the emergence of Omicron BA.1.

**Importance:** Our results highlight the plasticity of SARS-CoV-2 spike to generate highly fusogenic and cytopathic strains with the causative mutations being uncharacterized in previous variants. We describe mechanisms regulating the formation of syncytia and the subsequent consequences in cell lines and a primary culture model, which are poorly understood.

## INTRODUCTION

Over the timespan of the COVID-19 pandemic, SARS-CoV-2 has been subjected to selection pressures, leading to emerging variants carrying their own repertoire of mutations and to temporal waves of epidemiological resurgence (1, 2). The most successful variants evolved to evade the immune response (3–8) displaying differing abilities to form syncytia in cell culture systems (9, 10). During the period of co-circulation, the disease severity of Omicron (BA.1) was reduced in comparison to Delta (11–13). Proposed explanations for this include background immunity, different tissue tropisms - with BA.1 preferentially replicating in the upper respiratory tract - and reduced cell-cell fusogenicity of BA.1 spike (14–17). Therefore, the mechanisms surrounding SARS-CoV-2 pathogenicity and Omicron’s attenuation are still debated.

SARS-CoV-2 fuses with the cell plasma membrane to transfer its genome into the cytoplasm and instigate replication. This process is initiated through the binding of the spike to its receptor angiotensin-converting enzyme 2 (ACE2) (18). Spike is comprised of two subunits, S1 and S2, separated by a polybasic furin cleavage site (FCS) cleaved during viral production in the trans-Golgi network. Certain mutations in spike, such as P681H/R, allow for this process to occur more readily, subsequently improving viral fusion (19, 20). During entry, spike is cleaved at the S2’ site by host proteases, mainly TMPRSS2 at the cell surface (21, 22) or cathepsins in endosomes (23, 24), to allow the conformational changes necessary to project the fusion peptide into the host membrane, leading to membrane fusion. Thus, a series of proteolytic events regulate SARS-CoV-2 entry and tropism prior to replication of the viral RNA.

The later stages of the SARS-CoV-2 replication cycle occur in the endoplasmic reticulum (ER) and Golgi network. Here, the host protein COPI binds to the spike cytoplasmic tail and traffics it to the packaging site of SARS-CoV-2 virions (25). However, sub-optimal spike binding to COPI results in leakage to the plasma membrane. Consequently, spike at the cell surface may interact with ACE2 on neighbouring cells leading to cell-cell fusion and syncytia formation (21, 26).

Histological studies of lung tissue from severe COVID-19 patients describe the presence of abnormal pneumocytes and large, multinucleated syncytia (27, 28). Syncytia are also observed in the lungs of long-COVID patients who eventually succumb to the disease up to 300 days after their last negative PCR test (29). Syncytia could thus facilitate persistent infection, as seen in RSV infection (30), or contribute to pathogenesis. Syncytia formation by SARS-CoV-2 has also been demonstrated in various cell lines and in human iPSC derived cardiomyocytes (9, 31). Spike mutations can impact fusogenicity. Notably, D215G and P681R/H increase fusion, with the latter promoting furin S1/S2 cleavage. Conversely, N856K and N969K in Omicron decrease fusogenicity (16, 24). Nevertheless, the role of syncytia in SARS-CoV-2 replication and the impact on disease severity has yet to be fully explored.

Minor SARS-CoV-2 variants harbouring uncharacterized mutations represent opportunities in understanding certain viral processes. Variant B.1.640.1 was first identified in Republic of the Congo in April 2021 and found circulating in France in October 2021 (32). As of May 2023, 1107 genome sequences of B.1.640.1 are available on the GISAID database (33), 895 from France, with the most recent dating to January 2022. Here, we isolated a B.1.640.1 strain and investigated the humoral immune response, replication, fusogenicity and cytopathy of this variant.

## MATERIALS AND METHODS

### Plasmids

Human codon-optimised SARS-CoV-2 spikes (Alpha, Delta, BA.1, BA.5, B.1.640.1 and B.1.640.2) were produced *in silico* (GeneArt, Thermo Fisher Scientific). Spike sequences were then cloned into a phCMV backbone (GeneBank: AJ318514) using Gateway cloning (Thermo Fisher Scientific). Individual mutations (D614G, T859N, D936H) were produced through Q5 site-directed mutagenesis (NEB) in indicated backbones. Spike chimeras were generated through Gibson assembly of PCR generated fragments (NEB). Primers used for site directed mutagenesis and Gibson assembly are described in table 1. pQCXIP-Empty plasmid was used as previously described (34). All plasmids were sequenced by the Eurofins Genomics TubeSeq service with the primers listed in Table 1.

**Table 1.**
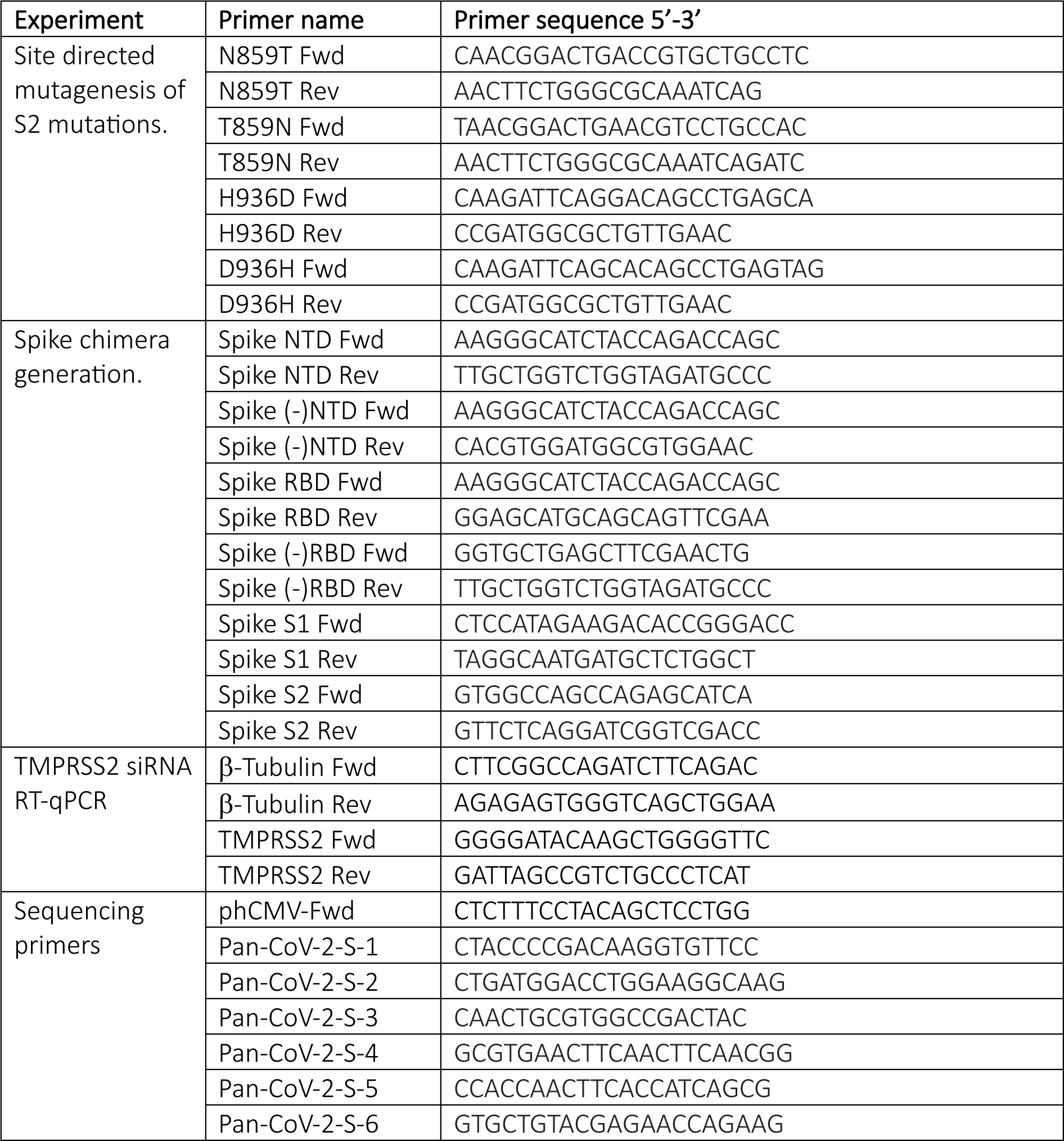
List of primers used throughout this study.

### Cells

HEK293T, Caco2/TC7, VeroE6, A549 cells, and derivatives were purchased from the ATCC or generously provided by fellow staff at Institut Pasteur. Cells were cultured in DMEM with 10% fetal bovine serum (FBS) and 1% penicillin/streptomycin (PS). GFP-split cells, expressing either GFP subunits 1-10 or GFP subunit 11, were generated through transduction of HEK293T and VeroE6 cells with respective pQCXIP-derived plasmids. Transduced cells were cultured with 1 μg/ml puromycin (InvivoGen). A549-ACE2 cells were generated through transduction of human ACE2 and cultured with 10 μg/ml blasticidin (21). All cells used in this study tested negative for mycoplasma.

### Cohorts

To study antibody neutralization of SARS-CoV-2 variants following infection or vaccination, a cohort from the prospective, monocentric, longitudinal, interventional cohort clinical study (ABCOVID) from Orléans was used. This study was commenced in August 2020 with the objective of studying the kinetics of COVID-19 antibodies in patients with confirmed SARS-CoV-2 infection (NCT04750720). A sub-study aimed to describe the kinetic of neutralizing antibodies after vaccination. The cohort is described in previous publications (3, 8, 35). To exclude individuals infected before vaccination, anti-N antibodies were measured upon sera collection. This study was approved by the Ile-de-France IV ethical committee. Written informed consent from the participants was collected and a questionnaire covering sociodemographic characteristics was completed.

### Origin of the B.1.640.1 strain

A nasopharyngeal swab collected from a patient tested positive for SARS-CoV-2 on November 2021, was sent to Hôpital Henri Mondor sequencing platform in the context of a nationwide survey. Briefly, private, and public diagnostic laboratories in France participated to the national SARS-CoV-2 genomic surveillance by providing a random subsampling of positive SARS CoV-2 samples to national sequencing platforms weekly (36). After sequencing, the leftover sample was used for viral isolation. The virus was isolated and amplified by one or two passages on VeroE6 cells as previously described (35).

### Virus sequencing

Viral RNA was extracted from the nasopharyngeal swab in viral transport medium. Sequencing was performed with the Illumina COVIDSeq Test (Illumina, San Diego, California), using 98-target multiplex amplifications along the full SARS-CoV-2 genome. The libraries were sequenced with NextSeq 500/550 High Output Kit v2.5 (75 Cycles) on a NextSeq 500 device (Illumina). The sequences were demultiplexed and assembled as full-length genomes using the DRAGEN COVIDSeq Test Pipeline on a local DRAGEN server (Illumina). The sample was identified as B.1.640.1 (hCoV-19/France/GES-HMN-21112100277/2021) before being submitted to the GISAID database (33), with the following ID: EPI_ISL_6470307. The sequence obtained by metagenomic sequencing (random hexamer cDNA generation and Nextera XT library preparation) after amplification was identical, and no contamination was detected.

### Other SARS-CoV-2 isolates

The D614G isolate was obtained through the European Virus Archive goes Global (Evag) platform and received from the National Reference Centre for Respiratory Viruses at Institut Pasteur. The Delta and BA.1 isolates were previously described (8). Strains were amplified and titrated on VeroE6 cells. Viral stock titres were calculated through TCID_50_ measurements. Viral stocks were diluted 1:10 in DMEM complete media and then serially diluted 10-fold for a total of 8 dilutions (10^-1^ to 10^-8^ dilution). Titrated virus was added to a 96-well plate containing 1.2×10^4^ VeroE6 cells per well. This was replicated six times per viral stock. Five days post-infection, cells were assessed for cytopathic effect using a light microscope. The lowest dilutions showing cytopathic effect were used to calculate the TCID_50_ of the viral stock. Manipulations of SARS-CoV-2 isolates were performed in a BSL-3 laboratory, under the guidelines of the risk prevention service of Institut Pasteur.

### GFP-Split fusion assay and video microscopy

To assess the fusion of the respective spike constructs, HEK293T-GFP1-10 cells (6×10^4^ cells per well) were transfected in suspension at 37°C using a shaking incubating at 750rpm for 30 min. The transfection mix was prepared using Lipofectamine 2000 (Thermo Fisher Scientific) with 50 ng DNA in a 1:10 ratio of SARS-CoV-2-S and pQCXIP-Empty respectively before being added to the cells. Following transfection, cells were washed and resuspended in DMEM 10% FBS. Transfected HEK293T cells were then co-cultured with either 1.5×10^4^ VeroE6-GFP11 cells or 1.5×10^4^ Caco2-GFP11 cells per well in a μClear black 96-well plate for 18 hours. Spike expression was assessed by staining with 1 μg/mL anti-S2 mAb (mAb10) targeting an unknown yet conserved epitope within the S2 subunit (8). For TMPRSS2 knockdown (KD) experiments, Caco-2-GFP11 cells were transfected with either control siRNA targeting luciferase (5ʹ-CGUACGCGGAAUACUUCGA-3ʹ) or siGENOME Humam TMPRSS2 SMARTpool (#7113 – Horizon Discovery) at 50 nM using Lipofectamine™ RNAiMAX (Thermo Fisher Scientific) for 48h. For TMPRSS2 inhibition, 10 μM camostat (CliniSciences, HY-13512) was added to Caco-2-GFP-11 cells 30 min prior to co-culturing with HEK293T-GFP1-10 cells. Hoechst 33342 was added to the media at a 1:10000 dilution. 18 hours post-transfection, images were acquired using the Opera Phenix High-Content Screening System (PerkinElmer) at 100x magnification. 20 images were acquired per well. Analysis was performed using Harmony High-Content Imaging and Analysis Software (PerkinElmer, HH17000012, v.5.0), including nuclei count and GFP area. For video microscopy, as described above, the cells were incubated (37°C, 5% CO_2_) inside the Mica Microhub system using the internal incubator (Leica microsystems). Videos were analysed using the complementary software (Leica microsystems).

### A549-ACE2 viral-infection, immunofluorescence, and video microscopy

6-hours prior to infection, 3×10^4^ A549-ACE2 cells were plated in a μClear black 96 well plate. Cells were then infected with MOI 0.1, 0.01, and 0.001 of SARS-CoV-2 virus. Cells were fixed using 4% PFA at 24-, 48-, and 72-hour time points and supernatant was collected for qRT-PCR and LDH analysis. Cells were then stained using 1 μg/mL of anti-spike mAb (mAb102, (35)) and 0.05% saponin in PBS, 1% BSA, 0.05% sodium azide for 30 min at room temperature. For the secondary antibody, cells were washed twice with PBS and stained as per the primary antibody with 1:600 goat anti-human IgG-FC alexa-647 antibody for 30 min then washed twice with PBS. Hoechst 33342 (Invitrogen) was added to the final PBS wash. Images and analyses were then performed as described above using the Opera Phenix High-Content Screening System. For video microscopy, A549-ACE2 cells were seeded in a µ-Dish 35 mm Quad (Ibidi) 6-hours prior to infection. Media was then replaced with DMEM 10% FBS with SARS-CoV-2 virus at an MOI of 0.1. Propidium iodide (PI) and Hoechst dyes were also added to the media. The microscopy was carried out using the BioStation video microscope, with six-fields acquired per chamber. Images were acquired every 10 mins over 36 hours. Video analysis and editing was performed using ImageJ software (Fiji).

### LDH activity assay

Cell supernatants were collected and stored in LDH storage buffer + Triton 10% as per the manufacturer’s instructions (Promega). Supernatants were inactivated for 2 hours at room temperature prior to usage outside the BSL-3. 50 μL of diluted supernatant was then added to 50 μL LDH substrate mix as described by Promega. LDH activity was recorded using the VICTOR3 multilabel plate reader (Perkin-Elmer).

### Human nasal epithelium culture, infection, and imaging

MucilAir^TM^, reconstructed human nasal epithelial cells (hNECs) that had been differentiated for 4 weeks prior to obtention, were cultured in 700 μL MucilAir^TM^ media on the basal side of the air/liquid interface (ALI) cultures and monitored for healthy cilia movements. One-hour prior to infection, mucus was removed from the apical side of the culture by washing the apical side with warm 200 μl MucilAir^TM^ media. Cells were then infected with equal virus titres in 100 μL MucilAir^TM^ media for 2 hours. Viral input was removed and stored at −80°C. Cells were then washed for 10 min at 37°C in warm PBS and then 20 min in 200 μL MucilAir^TM^ media for the day 0 recording. Washing with warm media was repeated every 24 hours for 96 hours. Every wash was subsequently centrifuged at 1500 rpm to remove cell debris and frozen at −80°C. After 96 hours, cells were fixed on the apical and basal sides with 4% PFA for 15 minutes. For imaging, fixed cells were stained intracellularly with rabbit anti-ZO1 (40-2200; Invitrogen), anti-SARS-CoV-2 N AlexaFluor-488 Dylight, as described in Planas et al., 2023, and rabbit anti-cleaved caspase-3 (D175; Cell Signaling Technology) and imaged using the LSM-700 confocal microscope (Zeiss).

### Flow cytometry

For surface staining, both primary and secondary antibodies were diluted with MACS buffer. Further information on the mAbs used in this study is listed in Table 2. Serum samples were diluted 1:300 and primary monoclonal antibodies were used at 1 μg/mL. Soluble-ACE2-human-IgG-FC was diluted to 10 μg/mL and serially diluted three-fold six times. Cells were mixed with 50 μL of primary antibody and incubated in the dark at 4°C. Cells were then washed in 100 μL PBS after staining. Goat anti-human FC AlexaFluor-647 was used as the secondary antibody at a dilution of 1:400. Cells were fixed in 4% PFA and acquired on an Attune NxT Flow Cytometer (Thermo Fisher). Data were analyzed using FlowJo software (BDBioSciences).

**Table 2.**
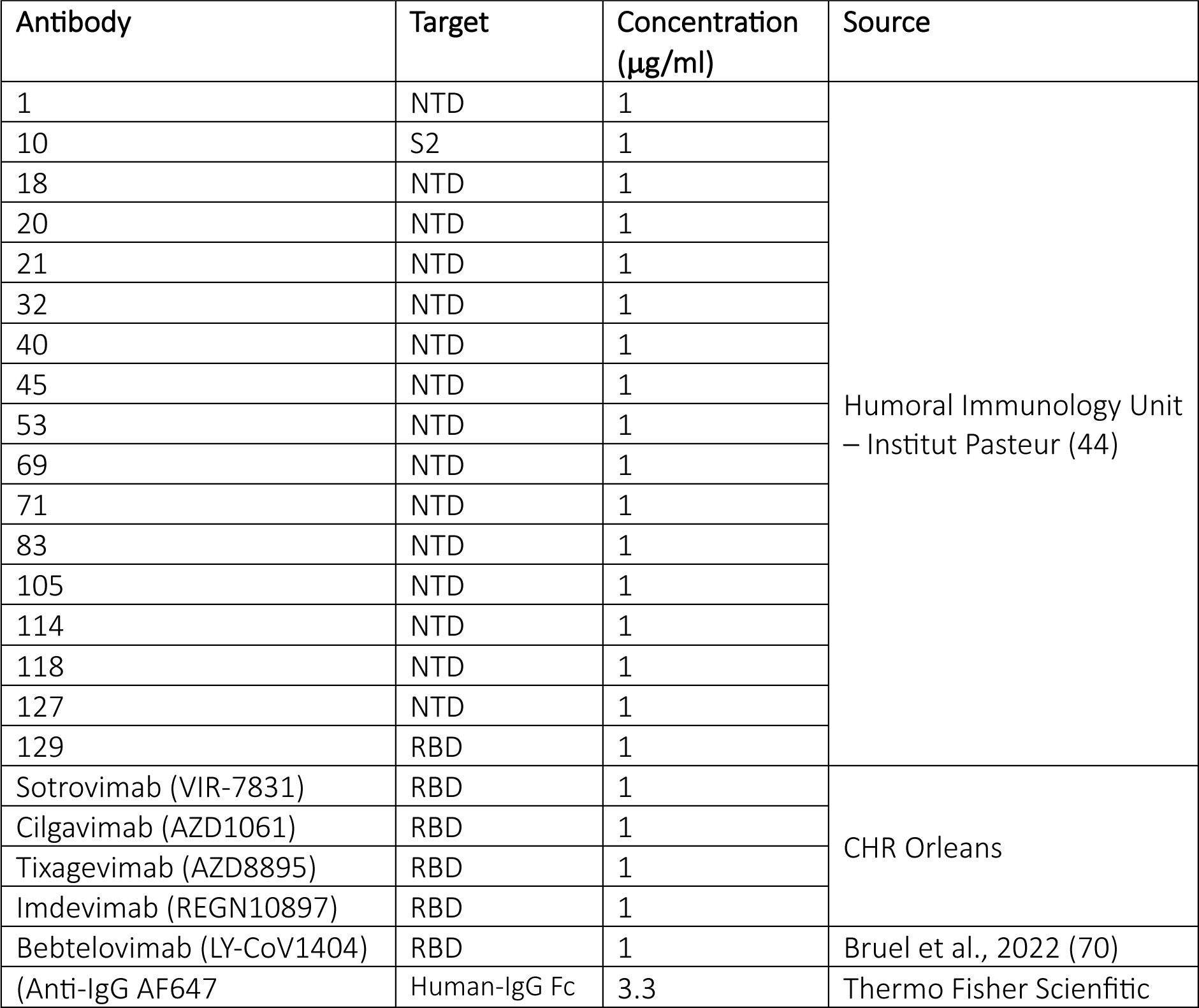
List of antibodies used throughout this study.

### S-Fuse neutralisation assay

U2OS-ACE2 GFP1-10 or GFP11, termed S-fuse cells, were mixed (1:1) and 8×10^3^ cells per well were plated in a μClear black 96 well plate. Heat inactivated serially diluted sera were incubated with the indicated SARS-CoV-2 viruses for 15 min at a starting dilution of 1:30 before being added to the cells. 18-hours post infection cells were fixed in 4% PFA and stained with Hoechst 33342 (Invitrogen). Cell images and GFP and nuclei quantification were performed using the Opera Phenix system as described above. Percentage neutralisation was calculated with the following formula: 100*(1-(x serum-x “non infected”)-(x no serum-x “non infected”)) with x = number of syncytia. Percentage neutralisation was then used to calculate the ED_50_ of each serum.

### RT-qPCR, RNA extraction, Reverse transcription

For quantification of viral RNA release, cell supernatants were collected and diluted 1:4 in H_2_O and then heat inactivated at 80°C for 20 min. 10 μM SARS-CoV-2 E-gene Forward (5’- ACAGGTACCTTAATAGTTAATAGCGT-3’) and Reverse (5’-ATATTGCAGCAGTACGCACACA-3’) primers were used with Luna Universal One-step RT-qPCR kit (New England Biolabs) and added to 1 μl supernatant (5μl total) in a 384-well plate. A standard curve was produced through a 1:10 serial dilution of EURM-019 ssRNA SARS-CoV-2 fragments for reference (European Commission). To evaluate Caco-2 TMPRSS2 KD, 5×10^5^ cells were lysed in RLT buffer (QIAGEN) with 10 μL of β-mercaptoethanol. RNA extraction and reverse transcription were performed using the RNeasy plus mini kit (QIAGEN) and Superscript II (Thermo Fisher Scientific), respectively, according to manufacturer’s instructions. RT-qPCR was carried out using iTaq universal SYBR green supermix (BioRad) and the primers listed in table 1. All RT-qPCR experiments were performed with a QuantStudio 6 Flex Real-Time PCR machine (Thermo Fisher Scientific).

### Scanning Electron Microscopy

MucilAir^TM^ samples were fixed with 4% PFA for 1h at RT to inactivate SARS-CoV-2 and stored in PBS at 4°C prior to preparation. Samples were then washed in 0.1 M cacodylate buffer and several times in water and processed by alternating incubations in 1% osmium tetroxide and 0.1 M thiocarbohydrazide (OTOTO). Samples were then dehydrated by incubating with increasing concentrations of ethanol. Samples were critical point dried and mounted on a stub for analysis. Analysis was performed by field emission scanning electron microscopy with a Jeol JSM6700F microscope operating at 3 kV.

### PyMOL

The SARS-CoV-2 spike structure (PDB: 6VXX) was imported in PyMOL. Mutations and residue proximites were calculated and annotated with the Mutagenesis and Measurement tools.

### Statistical analysis

Calculations were performed using Excel 365 (Microsoft). Figures and statistical analyses were conducted using GraphPad Prism 9. Statistical significance between different groups was calculated using the tests indicated in each figure legend. No statistical methods were used to predetermine sample size.

## RESULTS

### B.1.640.1 displays NTD antibody binding escape and reduced neutralisation sensitivity to Delta

The N-terminal domain (NTD) and Receptor Binding Domain (RBD) of spike are continuously subjected to immune selection pressures (2, 37). B.1.640.1 carries several unique mutations in these regions, including an unusually large NTD deletion of 9 amino-acids (Fig. 1A). To investigate their contribution to immune evasion, HEK293T cells expressing spikes from D614G, Delta, B.1.640.1, BA.1, and BA.5, were incubated with a set of anti-NTD monoclonal antibodies (mAbs) or anti-RBD therapeutic mAbs (Table 2). The therapeutic mAbs incubated at 1 μg/ml bound to all variants with the exceptions of Cilgavimab to B.1.640.1 and Tixagevimab to BA.5 (Fig.1B). Conversely, D614G and BA.5 showed a high level of binding to the anti-NTD mAbs, with 13/15 and 9/15 mAbs binding respectively. B.1.640.1 displayed the greatest reduction in anti-NTD binding with only 1 mAb retaining activity (Fig.1B).

**Fig 1.**
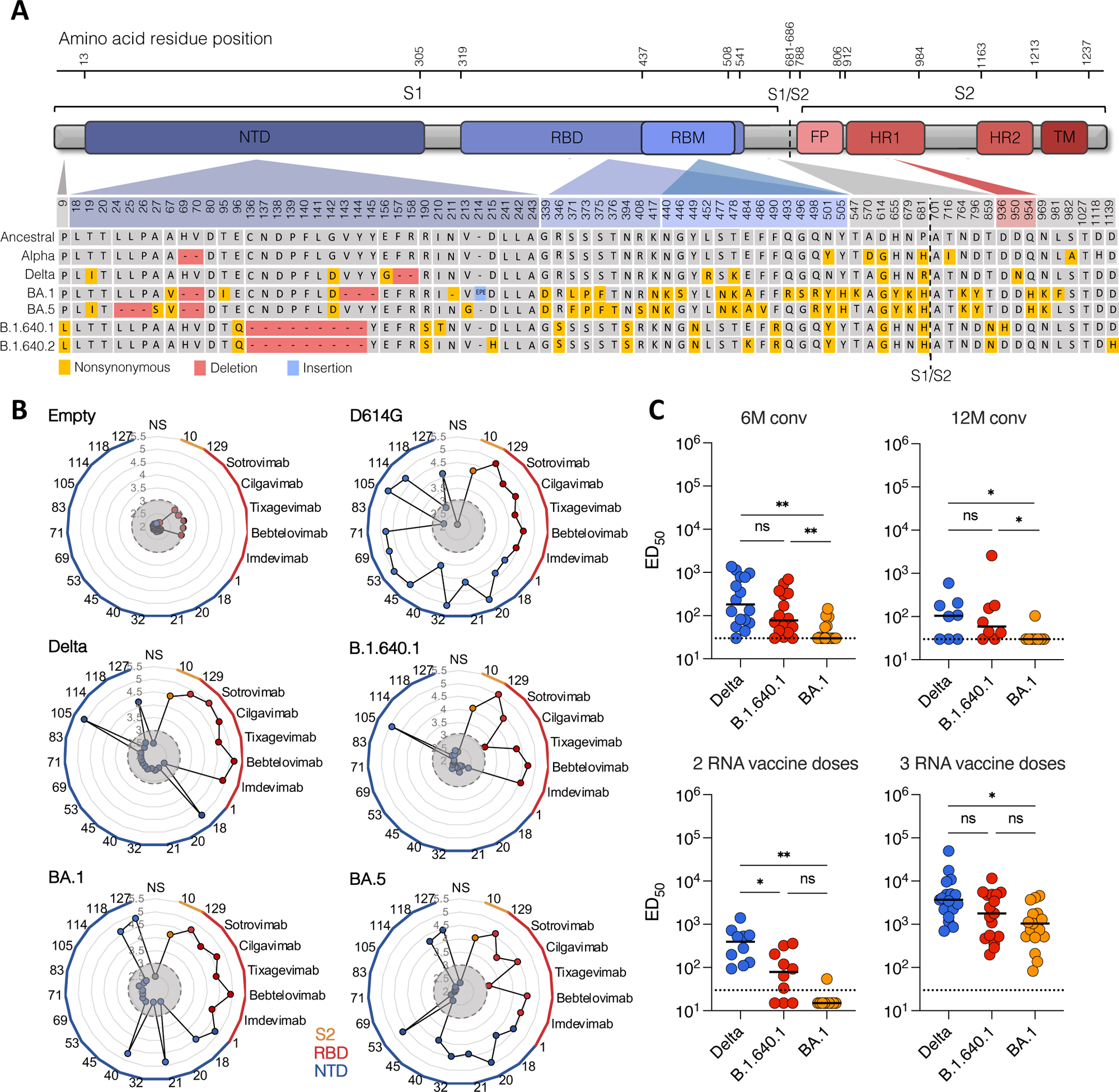
Monoclonal antibody binding and sera neutralization of B.1.640.1. A Schematic view of B.1.640.1 and B.1.640.2 sequences and their respective amino acid mutations as compared to the ancestral Wuhan sequence (NC_045512.2), Alpha, Delta, BA1, and BA5 variants. B Binding of a panel of mAbs targeting the S2, RBD, and NTD of spike of the respective variants with the y-axis showing log_10_(MFI). Grey circle indicates the limit of detection (MFI = log_10_(3)) C Neutralization activity of sera from convalescent individuals, 6- and 12-months after infection, and 1-month post second and third Pfizer RNA vaccines doses. Dotted line represents the limit of detection (ED_50_ =30). Solid black bars represent median values. Data points are a mean of two independent repeated experiments. Mann-Whitney tests were performed to compare the respective variants, \**P*<0.05, \*\**P*<0.001.

Next, we used a longitudinal cohort of sera from convalescent and vaccinated individuals established in the French city of Orléans, to test the sensitivity of B.1.640.1 to neutralizing antibodies. We isolated a B.1.640.1 strain from a swab of an infected individual from Hôpital Henri Mondor, Paris. Comparisons were drawn against BA.1 and to Delta which circulated at the same time as B.1.640.1. To measure neutralization, we calculated the 50% effective dilution (ED_50_) of each serum sample to each variant. B.1.640.1 showed a consistent reduction in neutralisation as compared to Delta with median ED_50_ values reduced by 2.3-fold and 1.7-fold, at month 6 (M6) and month 12 (M12), respectively (Fig.1C). As expected, BA.1 showed the greatest reduction in neutralization, with median ED_50_ values reducing 2.6-fold and 2.0-fold at M6 and M12, respectively, as compared to B.1.640.1 (Fig.1C).

Circulation of B.1.640.1 ceased in early 2022 at the time of Omicron emergence and concomitantly to the second and third RNA vaccine rollout in France. To investigate the sensitivity of B.1.640.1 to Pfizer vaccination, sera from individuals that received their 2^nd^ or 3^rd^ doses of vaccine was used to test viral neutralization. Here, Delta showed a 5.0-fold higher EC_50_ as compared to B.1.640.1 whereas BA.1 was minimally neutralized after two vaccine doses (Fig.1C). After three vaccine doses, B.1.640.1 displayed a 2.1-fold decrease and 1.7-fold increase in median EC_50_ over Delta and BA.1, respectively.

In all, B.1.640.1 spike showed anti-NTD mAb binding escape, with the consequences of this on the humoral response needing further investigation. Neutralization of the variant was consistently reduced as compared to Delta yet higher than BA.1. B.1.640.1 was efficiently neutralized after 3 doses of Pfizer vaccine.

### B.1.640.1 spike displays a high fusogenic phenotype

Viral fitness is influenced by factors such as immune evasion, tissue tropism, and replication capacity (17). To investigate the replication kinetics of B.1.640.1, compared to D614G, Delta and BA.1, we infected A549 human lung cell line expressing human ACE2 (referred to herein as A549-ACE2 cells) with a range of MOIs. Viral release was measured by RT-qPCR and the frequency of infected cells was quantified by spike immunostaining. At 72 hours post-infection (hpi) there was no significant difference in viral RNA release of B.1.640.1 or Delta as compared to D614G at MOI 0.01 and 0.1 (Fig. 2A). In accordance, no significant difference in the proportion of infected cells was observed between variants (Fig. S1A). BA.1 did not replicate in this cell line after 72 hpi (Fig. S1B).

**Fig 2.**
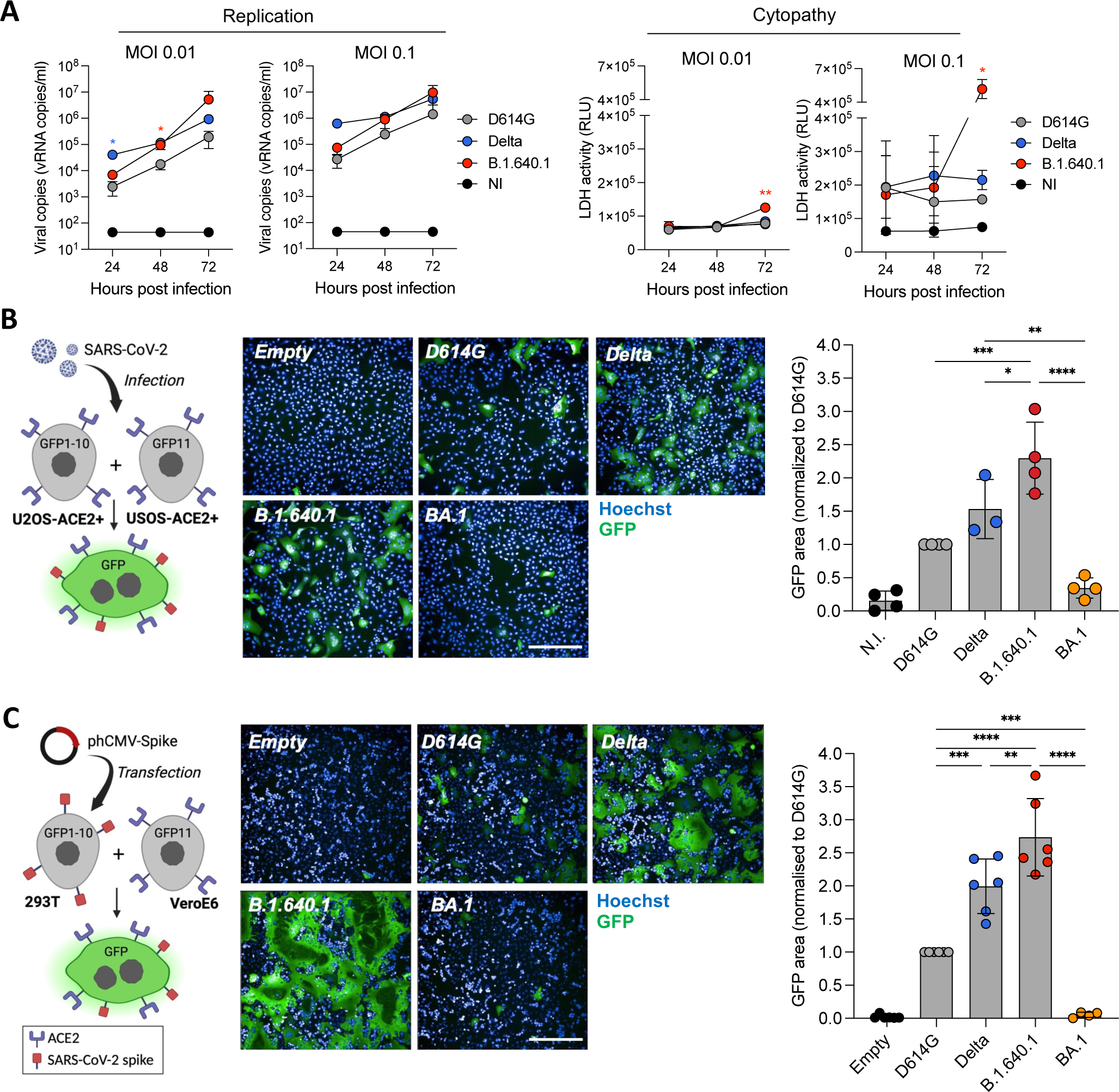
B.1.640.1 displays high spike fusogenicity and cytopathy in cell lines. A Replication kinetics of D614G, Delta, and B.1.640.1 in A549-ACE2 cells shown by quantification of the viral E protein gene in the cell supernatant at the respective timepoints (Left). Quantification of LDH release in A549-ACE2 infected cell supernatant through luciferase—based assay (Right). n=4. Two-way ANOVA tests were performed with Geisser-Greenhouse correction to compare Delta and B.1.640.1 to D614G, \**P*<0.05. B Schematic of S-fuse assay utilizing U2OS GFP-split cells to quantify viral fusion following infection (MOI = 0.01). Confocal microscopy images obtained at 200x magnification of the S-fuse system with respective variants at MOI 0.01. Quantification of GFP area was performed 18 hpi, data normalized to D614G fusion. C Schematic of HEK293T-spike GFP1-10 and VeroE6-GFP11 co-culture system. Fusion of transfected spike protein of respective variants through GFP area quantification 18 hours post-transfection. All images were obtained at 200x magnification. For B and C, ordinary one-way ANOVA tests were performed with Tukey’s multiple comparison test to compare D614G to respective variants, \**P*<0.05, \*\**P*<0.001, ***P<0.0001, ****P<0.00001. Error bars represent SD. Scale bars = 400 µm.

In contrast, microscopy analysis of B.1.640.1 infected cells revealed striking levels of cell-cell fusion at 48 hpi compared to other variants (Fig. S1C). To further investigate this phenomenon, we infected A549-ACE2 cells and monitored the cells using video microscopy over 48h. Video-microscopy analysis showed rapid fusion and syncytial formation in B.1.640.1 infected cells commencing 24 hpi, ultimately leading to large syncytia and cell death as detected by propidium iodide dye (Fig. S1D & Movie S1). D614G also induced syncytial formation but to a far lesser extent.

To confirm the high cytopathicity of B.1.640.1 observed by video microscopy, lactate dehydrogenase (LDH) activity was measured on cell supernatants to assess the level of plasma membrane damage in the infected cells. LDH activity peaked significantly at 72 hpi following B.1.640.1 infection at MOI 0.1 (5.2×10^5^ RLU and 1.6×10^5^ RLU for B.1.640.1 and D614G, respectively) indicating a surge in cytotoxicity (Fig. 2A). Infection of B.1.640.1 at MOI 0.1 also showed a significant reduction in nuclei number compared to Delta and D614G, with 2-6%, 17-33%, and 38-51%, fewer nuclei at 24h, 48h, and 72h respectively (Fig. S1E).

We then quantified cell-cell fusion using a GFP-split S-Fuse assay in U2OS-ACE2 cells (Fig. 2B), as described previously (9, 21, 35). B.1.640.1 infection at MOI 0.01 induced a 2.3-fold and 1.4-fold increase in cell-cell fusion as compared to D614G and Delta respectively (Fig. 2B).

Next, we investigated the fusogenicity of the variant spike proteins independently of viral replication. We transfected spikes into HEK293T cells expressing GFP1-10 subunits and incubated them with VeroE6 cells expressing GFP11 subunit for 18h (Fig. 2C). Spike expression on transfected HEK293T cells was similar across variants, as measured by anti-S2 mAb staining (Fig. S2B). As seen during infection, B.1.640.1 spike displayed the greatest fusogenicity as compared to other variants, with a 2.6-fold and a 1.3-fold increase over D614G and Delta respectively (Fig.2C). To further confirm the increased fusogenicity of B.1.640.1 spike, we transfected HEK293T GFP-Split cells with variant Spikes and ACE2 and monitored cell fusion by video microscopy over 18h (Fig. S2A & Movie S2). B.1.640.1 spike demonstrated a more rapid fusion kinetic across 18h of cell co-incubation.

Overall, B.1.640.1 showed a highly fusogenic phenotype and an increase cytopathy while having similar replication kinetics to the Delta in A549-ACE2 cells. In addition, B.1.640.1 spike alone displayed higher fusogenicity than D614G and Delta while being similarly expressed.

### B.1.640.1 infected human nasal epithelial cells (hNECs) form large syncytia

To investigate our findings in a physiologically relevant model, we infected MucilAir^TM^ primary human nasal epithelial cells (hNECs) in an air-liquid interface (ALI) culture with D614G, Delta, B.1.640.1, and BA.1 variants for 96 hours (Fig. 3A). This culture system is an effective tool to study SARS-CoV-2 infection (10, 14, 38). As previously described (5), BA.1 exhibited a replication advantage compared to Delta and D614G, as quantified by viral RNA release (Fig. 3B). Compared to BA.1, B.1.640.1 replication was delayed, with a 13-fold reduction in average RNA release after 24h of infection. B.1.640.1 replication kinetics closely resembled Delta after 48h of infection, with a 2.7-fold increase in peak Delta replication at 72 hpi (Fig. 3B). We next sought to examine if B.1.640.1 induced syncytia in this primary cell model. The cell cultures were fixed and stained 96 hpi when all variants had comparable viral RNA release (Fig. 3B). Due to the absence of fusion reporter systems, we assessed fusion through staining of Zonula Occludens 1 (ZO-1), a tight junction protein found a cell-cell borders, and phalloidin, binding to F-actin. ZO-1 and phalloidin staining revealed the presence of large, infected nucleoprotein-positive syncytia in the B.1.640.1 and Delta conditions (Fig. 3C, Fig. S3C). D614G induced smaller and rarer syncytia, however replicated to a far lesser extent. Conversely, BA.1 infection resulted in considerable destructuration of the epithelium and presence of small syncytia. However, its significantly higher replication in the ALI culture system may impact comparisons to the other variants. Delta and B.1.640.1 displayed similar replication kinetics, and both induced the formation of syncytia, with B.1.640.1 inducing large-rounded syncytia, confirmed through ZO-1 staining (Fig 3C, Fig.S3C). B.1.640.1 thus efficiently induces cell-cell fusion in primary hNEC ALI cultures with no replicative advantage as compared to Delta.

**Fig 3.**
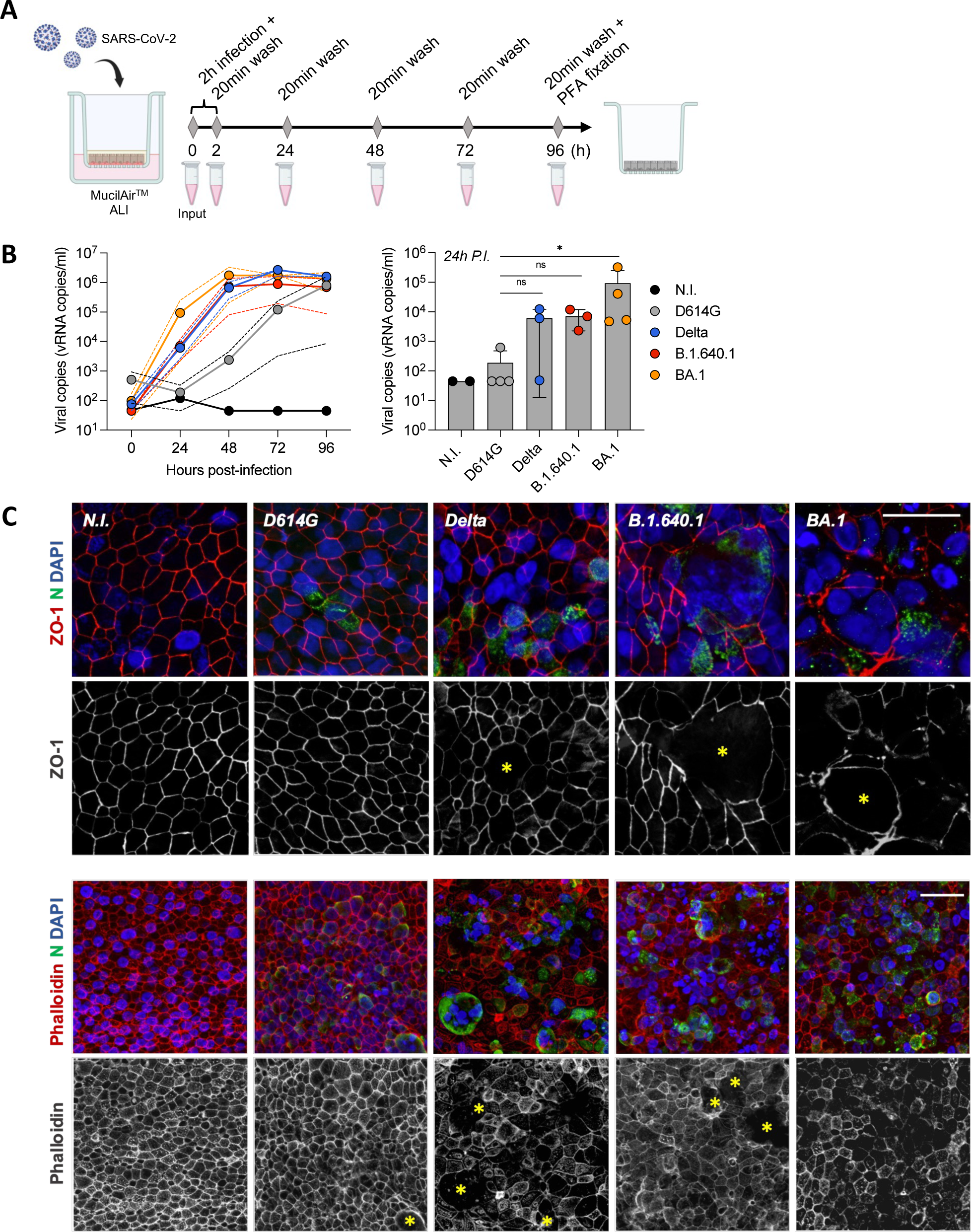
Apical replication and syncytia formation in hNEC infection. A Schematic hNEC ALI culture infection with SARS-CoV-2 variants over 96h followed by cell fixation. B Viral RNA release from apical side of infected hNECs as measured by RT-qPCR targeting the SARS-CoV-2 E protein RNA (n=3/4; Left). Apical viral RNA release/mL measured 24 hpi, bars represent mean values (Right). Dotted lines and error bars represent SD. Mann-Whitney tests were performed to compare the respective variants to D614G, \**P*<0.05. C Confocal immunofluorescence of hNECs 96 hpi with respective variants displaying syncytia formation through ZO-1, SARS-CoV-2 nucleoprotein, and DAPI staining. Scale bar = 20 µm.

### Elevated caspase-3 cleavage and LDH release during B.1.640.1 hNEC infection

Next, we investigated the cytopathy of B.1.640.1 in the primary cell model. We first quantified LDH release in the supernatant of infected cells. LDH release was detected at 72h and was significantly increased at 96 hpi for Delta and B.1.640.1, most notably for B.1.640.1 (Fig. 4A). BA.1 induced detectable but non-significant LDH while D614G did not induce detectable LDH release 96 hpi. As SARS-CoV-2 replication results in cytopathy (38–40) we normalised the LDH release to the levels of replication. Once normalised, B.1.640.1 infection increased LDH activity by 35% as compared to BA.1 following linear regression, suggesting B.1.640.1 is more cytopathic in primary nasal ALI cultures (Fig. 4A). To further investigate the mechanism of cell death, we examined the levels of cleaved caspase-3 induced by each variant. Caspase-3 undergoes cleavage following activation of the intrinsic or extrinsic cell death pathways; thus, its detection is a reliable marker for cell demise (41–43). Caspase-3 activation, assessed by immunostaining, was increased significantly by infection with B.1.640.1 as compared to D614G, Delta, and BA.1 (Fig. 4B and Fig. 4C). A similar but non-significant trend was seen in the replicate infection (Fig S3A). After normalizing for replication, B.1.640.1 infection resulted in a 2-fold increase in caspase-3 activation. Caspase-3 cleavage correlated significantly (P<0.005) with LDH release across all variants (Fig. 4D). In accordance with the cell line observations, B.1.640.1 displays elevated markers of cytopathy in hNEC ALI cultures.

**Fig 4.**
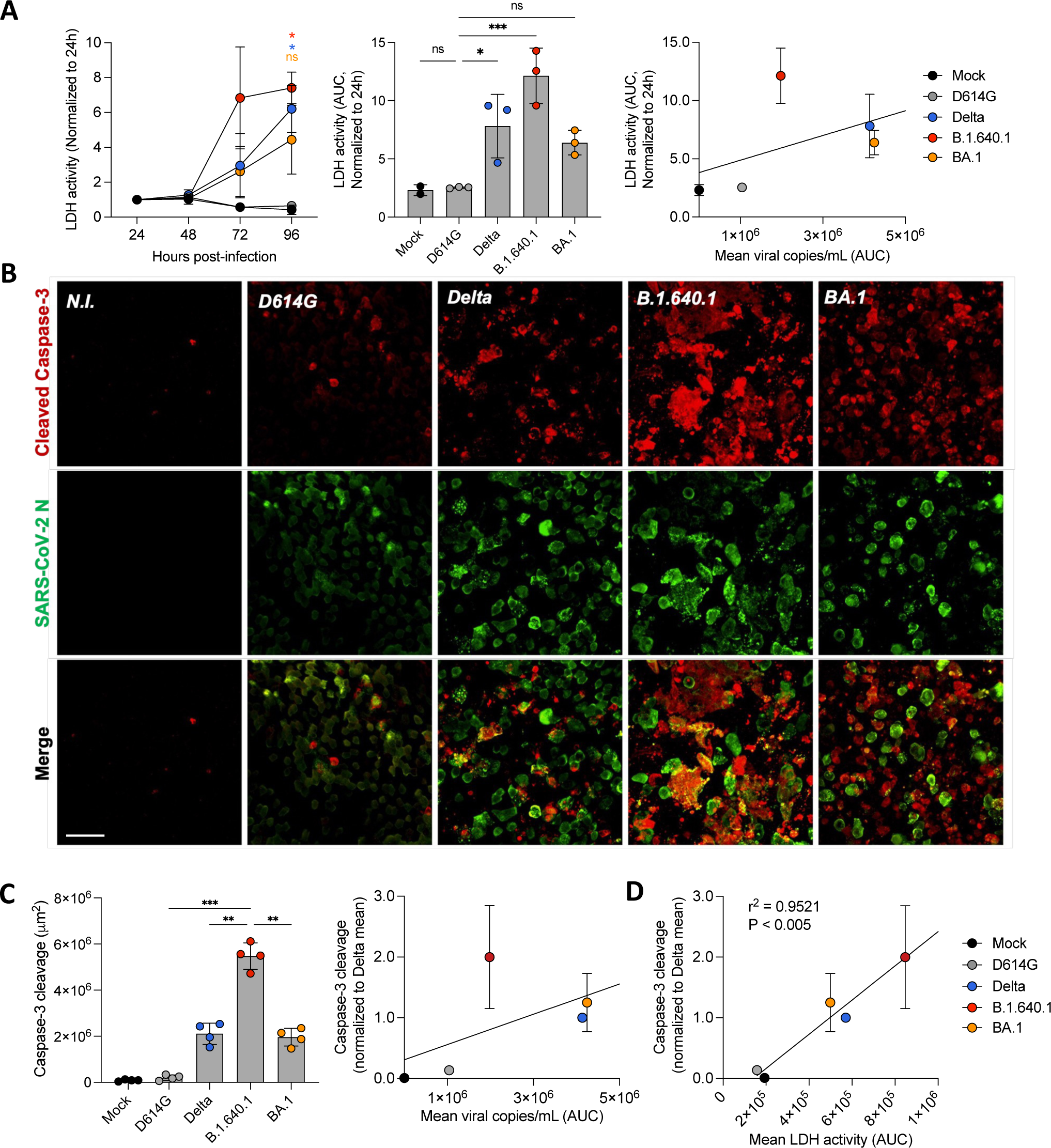
Cytopathic effects of hNEC infection with B.1.640.1. A LDH release from apical side of hNEC ALI culture over the time course of infection with respective SARS-CoV-2 variants (n=3/4; Left). Area under the curve (AUC) representation of LDH activity, bars represent mean values (Middle). Linear regression analysis of LDH release (AUC) as compared to viral copies/mL (AUC) from 96 hours of infection with respective SARS-CoV-2 variants (Right). B Immunofluorescence of hNECs stained for cleavage products of caspase-3 and SARS-CoV-2 nucleoprotein. Shown is one field of each variant. Scale bar = 40 µm. C Quantification of total area of cleavage products of caspase-3. Each data point represents one randomly assigned field from a single biological repeat (Left). An ordinary one-way ANOVA test was performed with Tukey’s multiple comparison test to compare D614G to respective variants, \**P*<0.05, \*\**P*<0.001. Linear regression analysis of caspase-3 cleavage from two biological repeats (Total number of fields = 8) normalized to Delta as compared to mean viral copies/mL (AUC) over 96 hours of infection (Right). D Linear regression analysis of caspase-3 cleavage normalized to Delta as compared to LDH activity (AUC) over 96 hours of infection. Error bars represent SD.

### S2 domain mutations T859N and D936H promote the high fusogenicity of B.1.640.1

We next explored the different mechanisms which could explain the increased fusogenicity of B.1.640.1 spike. We first checked the affinity of spike to ACE2 (Fig. 5A). Using a soluble ACE2-Fc binding assay, B.1.640.1 spike affinity was similar to that of D614G, suggesting that the increase in fusion is not due to differences in ACE2 affinity. Next, we investigated if B.1.640.1’s increased fusion can be explained by increased TMPRSS2 processing. TMPRSS2 acts synergistically with furin to process spike and enhance fusion (21, 22). We thus treated Caco-2-GFP11 cells with camostat, a TMPRSS2 inhibitor, and also performed siRNA knockdown of TMPRSS2 (Fig. 5B). Knockdown of TMPRSS2 was confirmed by RT-qPCR (Fig. S4B). Both camostat treated and siRNA treated Caco-2 cells were incubated with HEK293T-GFP1-10 cells expressing spike. This resulted in a 2.4-fold and 2.0-fold reduction in D614G and B.1.640.1 fusion, respectively (Fig. 5B). Nevertheless, B.1.640.1 spike maintained a 2-fold increase in fusion over D614G in TMPRSS2 inhibition or TMPRSS2 knockdown conditions. This result suggests that B.1.640.1 spike increases cell-cell fusion independently of its processing by TMPRSS2.

**Fig 5.**
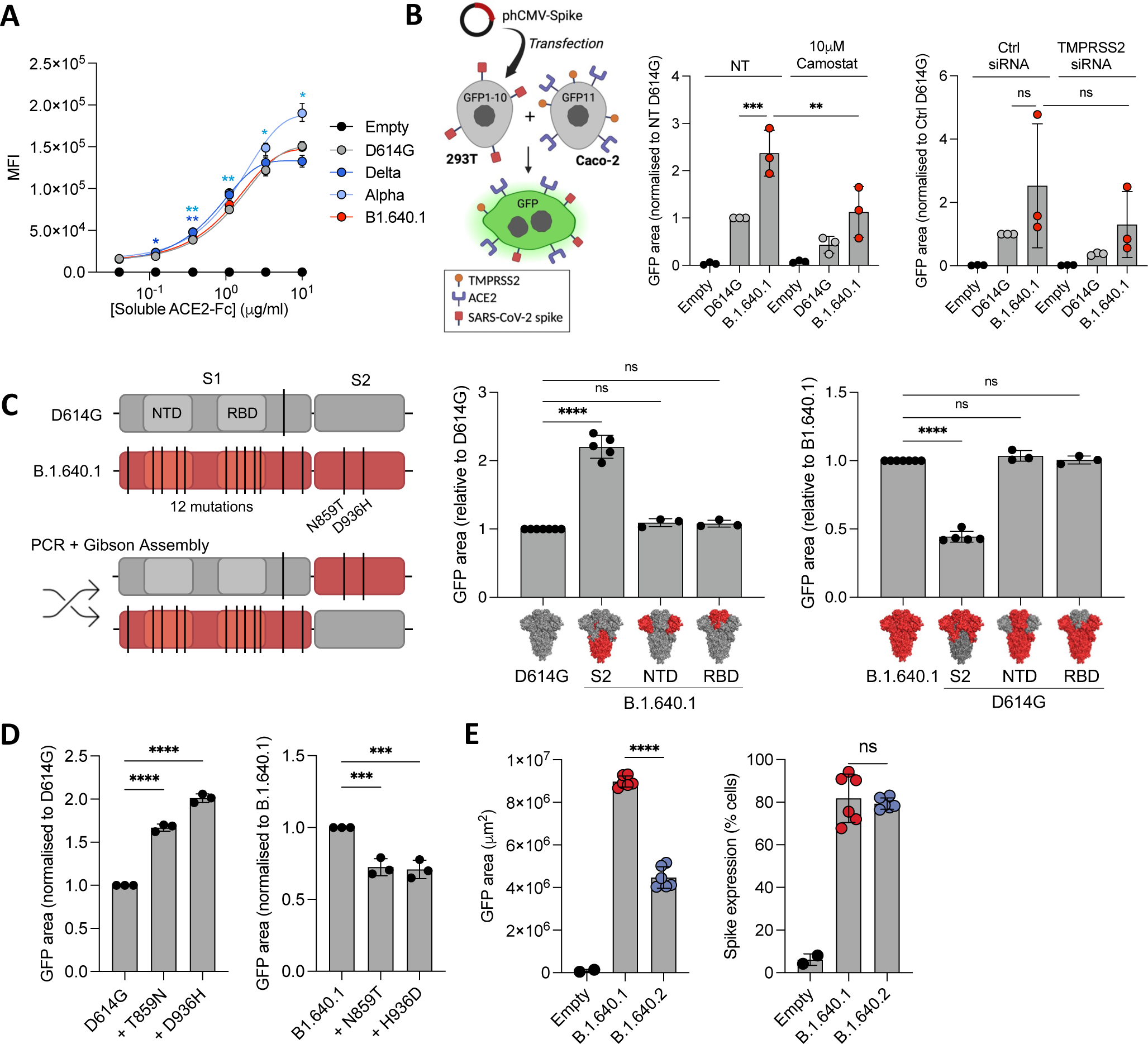
Spike S2 mutations of B.1.640.1 increase fusogenicity. A Staining of HEK293T cells transfected with spike and stained with a serial dilution of soluble human ACE2-Fc followed by staining with anti-human-Fc secondary antibody. A two-way ANOVA test was performed with Geisser-Greenhouse correction to compare D614G to the respective variants, \**P*<0.05, \*\**P*<0.001. B Schematic of HEK293T-spike GFP1-10 and Caco-2-GFP11 co-culture system (left). Fusion of D614G and B.1.640.1 spikes, quantified by GFP area, in the presence or absence of 10 μM camostat or TMPRSS2 siRNA knockdown (right). C Schematic of D614G and B.1.640.1 chimeric spike production through Gibson’s assembly (left). Fusion of D614G and B.1.640.1 spike chimeras in HEK293T-spike GFP1-10 and VeroE6-GFP11 co-culture system (right). D System is as described in C, B.1.640.1 and D614G spike fusion in the presence or absence of B.1.640.1 S2 mutations. E System is as described in C, B.1.640.1 and B.1.640.2 spike fusion. For B, C, D, E, ordinary one-way ANOVA tests were performed with Tukey’s multiple comparison test to compare D614G to respective variants, \**P*<0.05, \*\**P*<0.001, \*\*\**P*<0.0001, \*\*\*\**P*<0.00001. Error bars represent SD.

To identify the regions responsible for this phenotype, we generated chimeric spike plasmids by swapping the NTD, RBD, or S2 domains of D614G and B.1.640.1 spikes (Fig. 5C). We first confirmed that the chimeras were expressed at the surface at the same level (Fig. S4B). Next, using the HEK293T/VeroE6 GFP-split system, we observed that B.1.640.1 S2 approximately doubled D614G fusion while D614G S2 halved B.1.640.1 fusion while the NTD and RBD regions had no significant impact on fusion (Fig. 5C). Interestingly, only two mutations reside in the B.1.640.1 S2 domain, T859N and D936H. Using site directed mutagenesis, we introduced these mutations and their respective reversions individually into the D614G and B.1.640.1 spike backbones. Concurrent with the spike chimeras, both mutations significantly increased D614G fusion while the reversion mutations significantly reduced B.1.640.1 fusion (Fig. 5F) while being expressed similarly (Fig. S4C). Altogether, these findings reveal that the increased fusogenic property of B.1.640.1 is linked to the two S2 subunit mutations T859N and D936H.

### B.1.640.2 spike is less fusogenic than B.1.640.1

The sister variant of B.1.640.1, B.1.640.2, circulated in France over a similar timespan. The two variants differ by five mutations in the spike protein (Fig. 1A). Notably, the D936H HR1 mutation is not present in B.1.640.2. Consistent with this, fusion of B.1.640.2 spike was reduced by 2-fold in comparison to B.1.640.1 (Fig. 5E) while being expressed similarly (Fig. 5 E). This is a greater reduction than the single point mutation alone, suggesting that other mutations within B.1.640.2 reduce its fusogenic potential.

## DISCUSSION

We describe the phenotypic properties of the SARS-CoV-2 B.1.640.1 variant which transiently circulated in Europe and Africa in 2021-2022. Analysis of a panel of mAbs identified a loss of anti-NTD binding to this variant. The role of the NTD in neutralizing and non-neutralizing antibody functions is of current interest (44–46). A region termed the NTD antigenic supersite is a favourable target for potently neutralizing antibodies (2, 47). The supersite comprises of three loops, N1 (residues 14-26), N3 (residues 141-156), and N5 (residues 246-260). Deletion or mutation of these residues and proximal residues, such as C136Y, G142D, Del144, greatly reduces neutralization by mAbs by changing loop conformations and antibody epitopes (47). The epitopes of the NTD mAbs we tested are unknown. It is likely that Del136-144 in B.1.640.1 NTD results in alteration to the antibody supersite and thus loss of antibody recognition, but this requires further structural analysis.

B.1.640.1 spike demonstrates a plasticity of the NTD to undergo large levels of mutation while maintaining the protein functions. Supersite deletions are seen during persistent infections of immunocompromised individuals (48, 49). Additionally, the NTD is a region of interest for non-neutralizing antibody (nnAb) functions such as antibody-dependent cell-cytotoxicity (ADCC) and complement-dependent cytotoxicity (CDC) (44, 46, 50). Future studies would be of interest to reveal if the B.1.640.1 NTD and Del136-144 arose as a mechanism of nnAb evasion.

Our mAb binding analysis revealed the loss of Cilgavimab (AZD1061) binding to B.1.640.1, whereas other anti-RBD mBA maintained their binding capacities. A similar loss of binding was seen in recent Omicron subvariants BA.2.75.2 and BQ.1.1, all containing spike mutation R346T, now known to cause the loss of binding (3, 51, 52). As such, Cilgavimab is no longer recommend for therapeutic use. B.1.640.1 carries a R346S substitution, demonstrating that convergent mutations with clinical relevance have circulated in older variants.

Emergence and dominance of variants of concern are facilitated by many viral and host factors. The success of the initial Omicron variant, BA.1, was mainly due to its considerable evasion of neutralising antibodies (8, 53–56), among other factors. B.1.640.1 displayed a consistent reduction in neutralization as compared to Delta by sera from convalescent or vaccinated individuals. Such a reduction may have allowed the spread of B.1.640.1 in France while Delta was prevalent. BA.1 showed a significant reduction in neutralization compared to Delta in all conditions and B.1.640.1 in sera from infected individuals. Additionally, the E484K mutation is important for immune evasion but is absent in B.1.640.1 (57). Therefore, while B.1.640.1 was able to circulate within France, its disappearance came with the rollout of the third SARS-CoV-2 vaccine dose and the emergence of BA.1.

We report the ability of B.1.640.1 to induce large syncytia in our culture systems. Syncytia are observed in histological studies of COVID-19 patients’ lungs however their role in pathogenesis is still debated and their presence during mild disease remains unclear (28, 29, 58). We found B.1.640.1 elevated LDH release and activation of caspase-3 beyond that of the other variants. Cell death pathways are tightly regulated through caspase activation (59). Caspase-8, a master regulator that cleaves caspase-3/7, is upregulated in SARS-CoV-2 infected lung epithelial cells (60, 61). The replication of SARS-CoV-2 is sufficient to cause cytopathic effects in airway epithelial cells (38, 40, 62). This is consistent with the ciliopathy observed in the primary nasal epithelium cultures (Fig. S3B). Nevertheless, B.1.640.1 caused greater LDH release and caspase-3 cleavage than the other variants when normalized for replication. In accordance, SARS-CoV-2-spike-induced syncytia formation in HeLa cells causes activation of caspase-3 and pyroptosis, an inflammatory cell death pathway (41). Cell-cell fusion induced by FAST fusion protein of reoviruses enhances viral pathogenicity (63). Together, our results suggest that the highly fusogenic property of B.1.640.1 is responsible for its increased cytopathy.

We show that T859N and D936H within the B.1.640.1 S2 subunit are independently and additively responsible for high intrinsic fusogenicity of spike. Both mutations have continuously circulated at a low level (<0.05% of sequences), with T859N most notably being in the Lambda variant, and D936H being in a small proportion of omicron subvariants (covSPECTRUM; GISAID). The absence of these mutations in the Delta variant also suggests enhanced fusogenicity can arise through differing mutations. Additionally, B.1.640.1 spike fusion was increased in the presence of TMPRSS2. Similar levels of inhibition to D614G suggest a preferential cell surface entry route for B.1.640.1, consistent with variants prior to Omicron BA.1 (64). The S2 subunit contains the membrane fusion machinery including the hydrophobic fusion peptide (FP) and the heptad repeat regions 1 and 2 (HR1 and HR2). Mutations can alter this process through different mechanisms. For example, N856K present in BA.1 introduces a charged lysine residue that forms a salt-bridge with D568 which reduces shedding of the S1 subunit and fusion (16). *In silico* analysis reveals that residue 859 resides in the vicinity of residue 614 within S1 (Fig. S4D). It would be of interest to study any potential interactions between residue 859 and the S1 subunit and if subsequent mutations impact this interaction. HR1 and HR2 within S2 form a six-helical bundle to bring the membranes together during fusion (65). D936 within HR1 forms a salt bridge with the positively charged R1185 residue of HR2 (66). Thus, the introduction of H936 may also impact the conformational changes spike undergoes during fusion. Deep mutational scanning may also reveal how these mutations alter the effect of other spike mutations to understand the evolutionary constraints of fusogenicity on Spike (67).

*In silico* analyses suggest B.1.640.2 to have greater infectivity than B.1.640.1 (Pascarella et al., 2022). We previously showed the E484K mutation, present in B.1.640.2 but not B.1.640.1, significantly reduces fusion (9) and contributes to immune evasion (57). Together with the absence of D936H, this explains the 2-fold reduction in fusogenicity of B.1.640.2. Further investigation into the evolution of the B.1.640 lineage would be relevant to determine the relationship between B.1.640.1 and .2.

We acknowledge that this study contains limitations. Firstly, the low number of infections caused by B.1.640.1 limits the availability of clinical information on this variant. Further research may involve *in vivo* models, such as Syrian Hamsters or mice (38, 69), to assess B.1.640.1 pathogenicity. Additionally, we did not explore the role of other mutations outside of the spike in B.1.640.1 replication and cytopathy which could be studied through reverse genetic approaches. Further research to investigate the role of cytopathy in viral evolution would be relevant in understanding the evolution of SARS-CoV-2.

Overall, the data presented here shows a now supplanted SARS-CoV-2 variant B.1.640.1 displaying a highly fusogenic phenotype. Analysis of the humoral response provides insight into how this variant, like many others, were displaced upon the emergence of BA.1. The unusually high fusogenic activity of B.1.640.1, linked to an increased cytopathy, provides insight into consequences of SARS-CoV-2 cell-cell fusion.

## ACKNOWLEDGEMENTS

The authors thank the patients who contributed to this study, the members of the Virus and Immunity Unit (Institut Pasteur) and other teams for discussion and help, Nathalie Aulner and the staff at the UtechS Photonic BioImaging (UPBI) core facility (Institut Pasteur). W.B. was supported by a stipend from the Pasteur - Paris University (PPU) International PhD Program.

## FUNDING

Work in O.S. lab is funded by Institut Pasteur, Urgence COVID-19 Fundraising Campaign of Institut Pasteur, Fondation pour la Recherche Médicale (FRM), ANRS-MIE, the Vaccine Research Institute (ANR-10-LABX-77), Labex IBEID (ANR-10-LABX-62-IBEID), ANR / FRM Flash Covid PROTEO-SARS-CoV-2, ANR Coronamito, HERA European funding, Sanofi and IDISCOVR. The E.S.-L. laboratory receives funding from the Institut Pasteur, the INCEPTION program (Investissements d’Avenir grant ANR-16-CONV-0005) and NIH PICREID program (Award number U01AI151758). The Opera system was co-funded by Institut Pasteur and the Région ile de France (DIM1Health). The funders of this study had no role in study design, data collection, analysis, and interpretation, or writing of the article.

## AUTHOR CONTRIBUTIONS

Conceptualization: W.B., J.B., O.S., T.B., V.M.

Methodology: W.B., J.B., O.S., T.B., V.M., D.P., F.G-B., F.P., M.H., I.S.

Investigation: W.B., J.B., O.S., T.B., V.M., D.P.

Data Collection and Analysis: W.B., J.B., V.M., T.B., D.P., M.H.

Cohort design: L.H., T.P.

Resources: E.S-L., H.M., M.P., C.P., J-M.P., M.N., C.R., S.F., L.H., T.P.

Manuscript Writing and Editing: W.B., T.B., O.S., J.B.

## DECLARATION OF INTERESTS

The authors declare no competing interests.

---

**Supplementary figure 1. Cytopathic effects of B.1.640.1 on A549-ACE2 cells.**

**A** Area of spike-positive A549-ACE2 infected cells from respective variants over 72 hours.

**B** Replication kinetics of BA.1 in A549-ACE2 cells shown by quantification of the viral E protein gene in the cell supernatant at the respective timepoints.

**C** Confocal immunostaining of D614G, Delta, and B.1.640.1 infected A549-ACE2 cells 48 hours post-infection. Yellow arrows indicate the presence of syncytia. Scale bar = 200 µm.

**D** Video microscopy analysis of A549-ACE2 infected cells at indicated timepoints from Movie 1. Yellow borders highlight syncytia. Scale bar = 160 µm.

**E** Nuclei count following infection of A549-ACE2 cells at the indicated timepoints with the respective variants.

For **A** and **E**, two-way ANOVA tests were performed with Geisser-Greenhouse correction to compare Delta and B.1.640.1 to D614G, \**P*<0.05. Error bars represent SD.

**Supplementary figure 2. Kinetics of spike fusogenicity.**

**A** Fusion of variant spike proteins in HEK293T GFP1-10 cells, transfected with spike, and VeroE6-GFP11 co-culture system. Snapshots are shown from video microscopy at indicated timepoints. GFP area quantification was performed using six randomly assigned fields per variant. Scale bar = 200 µm.

**B** Number of syncytia produced by fusion of respective variant spike in HEK293T-spike GFP1-10 and VeroE6-GFP11 co-culture (Left). Staining of the HEK293T-spike cells with an anti-S2 mAb (Right). Spike expression was measured by percentage positive cells as compared to empty plasmid transfection. Ordinary one-way ANOVA tests were performed (on the final timepoint of **A**) with Tukey’s multiple comparison test to compare D614G to respective variants, \**P*<0.05, \*\**P*<0.001, \*\*\**P*<0.0001.

**Supplementary figure 3. Fluorescence microscopy and SEM of infected primary hNEC ALI cultures.**

**A** Caspase-3 cleavage 96 hpi from a second biological replicate of a hNEC infection. Each data point represents a randomly assigned field. Ordinary one-way ANOVA tests were performed with Tukey’s multiple comparison test to compare D614G to respective variants, \**P*<0.05, \*\**P*<0.001, \*\*\**P*<0.0001.

**B** SEM on SARS-CoV-2 infected primary nasal epithelia fixed 96 hours post-infection with the respective variants. Green colouration was added post-acquisition to highlight cilia. Scale bar = 200 µm.

**C** Confocal immunofluorescence of hNECs 96 hpi with respective variants displaying syncytia formation (yellow asterisks) through ZO-1, phalloidin, SARS-CoV-2 nucleoprotein, and DAPI staining. Upper scale bar = 20 µm. Lower scale bar = 40 µm.

**Supplementary figure 4.** TMPRSS2 knockdown, expression of spike chimeras and mutants, and *in silico* spike mutation analysis.

**A** Total RNA was extracted from Caco-2/TC7 cells treated with TMPRSS2 siRNA or siCTRL and RT-qPCR was performed. Data were normalized to *β-Tubulin* levels. Relative mRNA expression normalized to siCtrl condition (2−ΔΔCT) was plotted.

**B** Spike expression analysis using flow cytometry of HEK293T cells transfected with the respected spike chimeras for cell-cell fusion assays.

**C** As with B but for HEK293T cells transfected with respective spikes carrying mutations at residues 859 and 936.

**D** Structural analysis of residue 859 and it’s close proximity to residue 614 in the S1 subunit (PyMOL. Schrödinger).

For **B** and **C**, ordinary one-way ANOVA tests were performed with Tukey’s multiple comparison test to expression of respective spikes. Error bars represent SD.

**Supplementary movie 1.** Video-microscopy of A549-ACE2 cells infected with indicated SARS-CoV-2 variants at MOI 0.1. Scale bar = 80 µm.

**Supplementary movie 2.** Video-microscopy of HEK293T-GFP1-10 cells transfected with indicated SARS-CoV-2 variant spikes and co-cultured with VeroE6-GFP11 cells over 18-hours. Scale bar = 160 µm.

